# Peptide aggregation-induced immunogenic cell death in a breast cancer spheroid model

**DOI:** 10.1101/2023.10.31.565012

**Authors:** Gokhan Gunay, Katelyn N. Maier, Seren Hamsici, Filipa Carvalho, Tristan A. Timog, Handan Acar

## Abstract

Utilizing multicellular aggregates (spheroids) for in vitro cancer research offers a physiologically relevant model that closely mirrors the intricate tumor microenvironment, capturing properties of solid tumors such as cell interactions and drug resistance. In this research, we investigated the Peptide-Aggregation Induced Immunogenic Response (PAIIR), an innovative method employing engineered peptides we designed specifically to induce immunogenic cell death (ICD). We contrasted PAIIR-induced ICD with standard ICD and non-ICD inducer chemotherapeutics within the context of three-dimensional breast cancer tumor spheroids. Our findings reveal that PAIIR outperforms traditional chemotherapeutics in its efficacy to stimulate ICD. This is marked by the release of key damage-associated molecular patterns (DAMPs), which bolster the phagocytic clearance of dying cancer cells by dendritic cells (DCs) and, in turn, activate powerful anti-tumor immune responses. Additionally, we observed that PAIIR results in elevated dendritic cell activation and increased antitumor cytokine presence. This study not only showcases the utility of tumor spheroids for efficient high-throughput screening but also emphasizes PAIIR’s potential as a formidable immunotherapeutic strategy against breast cancer, setting the stage for deeper exploration and potential clinical implementation.

## 1. Introduction

In 2023, the U.S. is expected to see 300,000 new breast cancer diagnoses and 40,000 deaths.^1^ Immunotherapies, while promising for 15-20% of patients, leave many facing challenges like autoimmunity or distant relapse if their cancers remain unresponsive.^1–5^ Triple-negative breast cancer (TNBC) is particularly worrisome due to its resistance to chemotherapies.^1^

Establishing the optimal biological dose (OBD), the lowest dose providing the highest rate of efficacy while being safely administered, is essential for clinical trials. However, the factors determining the efficacy of therapies, such as tumor-infiltrating lymphocytes and immune checkpoints, are often observed when the detrimental effects have already become irreversible, making identifying the OBD challenging.^2,6^ This challenge accentuates the need for robust preclinical assessments. Clinical trial data across multiple cancer types pinpoints immunogenic cell death (ICD) as a cornerstone in immunotherapy’s efficacy. During ICD, distressed cancer cells release damage-associated molecular patterns (DAMPs), influencing tumor progression.^2,7–12^ Inducing cell death has been the primary goal of cancer treatment strategies for decades. However, emerging results show that the type of cell death dictates tumor progression. For example, induction of apoptosis within the tumor microenvironment (TME) leads to increased proliferation of cancer cells, reduced inflammation, and suppressed immune system, all of which increase cancer progression.^13^ Conversely, the release or surface expression of DAMPs in ICD can engage the immune system.^10^ Moreover, ICD leads to the release of tumor-associated antigens and promotes the initiation of antigen-specific adaptive immunity by activating dendritic cells (DCs)10,^14^. Different techniques induce ICD and DAMP release, such as chemotherapeutics^15^, photothermal therapy^16^, radiation therapy^17^, and recently introduced by our group, the peptide-aggregation model (PAIIR)^18^.

To induce ICD, we have developed a peptide-aggregation induced immunogenic response (PAIIR) method, which uses engineered peptides made of natural amino acids^19,20^. Inspired by the structures of transmembrane pore-forming proteins^21–23^, we utilized our molecular framework^19^ and designed a pair of oppositely charged peptides [II] that penetrate cells, aggregate, and induce stress and ICD^18,24^. Accumulation of misfolded proteins and their aggregates in the cells cause stress in the endoplasmic reticulum, and depending on the severity of the stress, it may stimulate eukaryote initiation factor 2 alpha (eIF2-α) phosphorylation and calreticulin exposure, which are hallmark biomarkers of ICD^25^. We demonstrated that the induction of ICD by PAIIR involves the phosphorylation of eIF2 and the exposure of calreticulin^26^. Furthermore, we observed that PAIIR can trigger ICD and the release of DAMPs in various cell lines, leading to an enhanced immune response against the influenza vaccination model^18^. Notably, our research also unveiled that using albumin as a modulator can control both peptide aggregation on the cell membrane and their subsequent effects on ICD and DAMP release **(Figure 1)**^24^.

**Figure 1.**
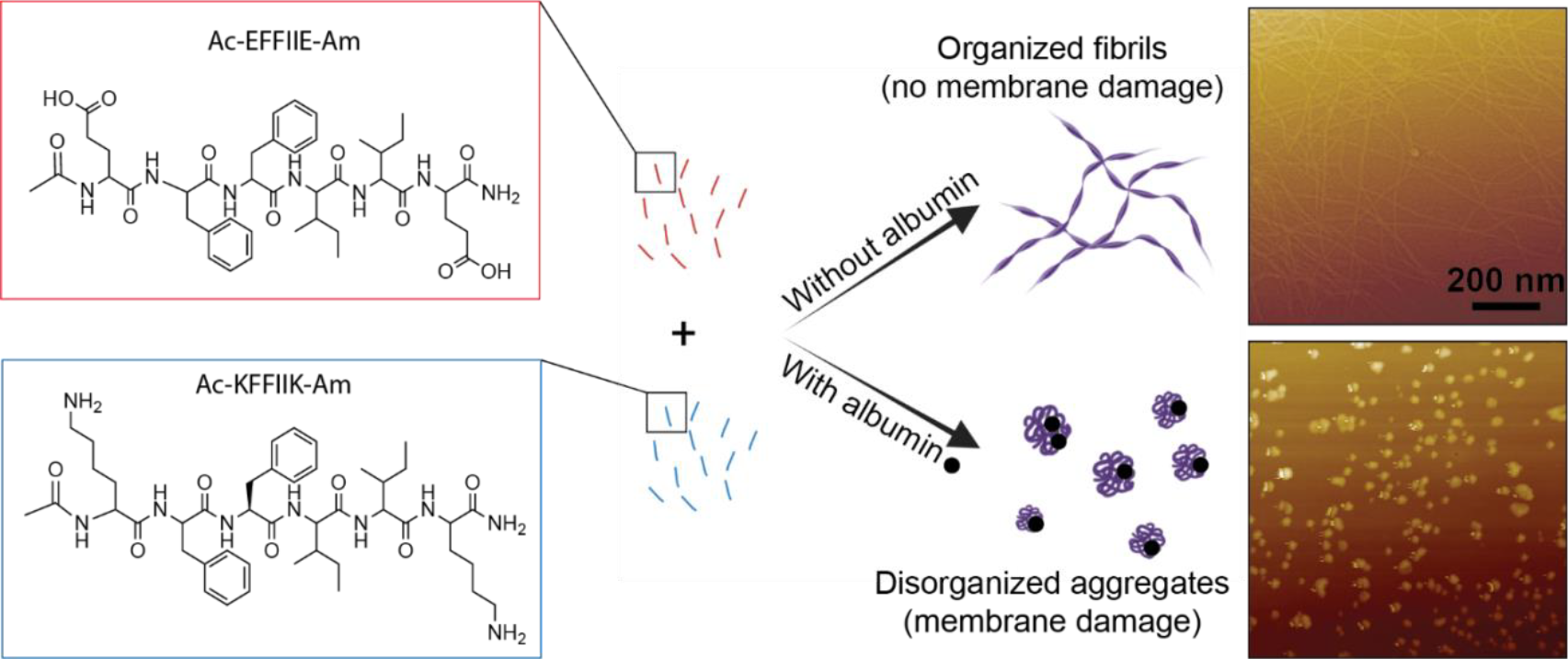
Co-assembly of oppositely charged peptides. [II] (EFFIIE--KFFIIK) self-assembles into fibrils (no membrane damage) in the absence of albumin and forms disorganized aggregates that induce membrane damage in the presence of albumin.

In preclinical studies, ICD and DAMP release via small drugs has been predominantly studied in monolayer cultures. Yet, these fail to mimic the complex nature of in vivo tumors, and while animal models offer a closer match, their cost and time-intensive nature pose significant barriers^27^. Utilizing multicellular aggregates of cancer cells, tumor spheroids, represent a bridge between in vitro and in vivo models to provide a physiologically relevant, cost-effective, and high-throughput model.

Cultivating cells in a three-dimensional (3D) format to form compact spheroids can better represent key tumor characteristics found in vivo compared to conventional two-dimensional (2D) cell cultures. Spheroid models offer physiological relevance to in vivo solid tumors in cell-to-cell interactions, cell-to-extracellular matrix interactions, and zone formation due to reduced nutrient and oxygen diffusion^28^. Importantly, gene expression shifts in spheroids versus monolayers can significantly alter the biological outcomes of cell death-inducing chemotherapeutics^29^.

For example, a recent study showed the changes in caspase expression upon paclitaxel and doxorubicin that resulted in the anti-apoptotic stage of tumor cells in breast cancer spheroids. Increased autophagy in breast cancer spheroids reduces the activity of trastuzumab, making spheroids resistant^30^. Another study showed that spheroids were resistant to ferroptosis (a form of regulated necrosis) induction compared with their monolayer counterparts^31^. These efforts highlight the importance of using physiologically relevant models to study and pre-select new therapeutics.

Induction of ICD within the tumor microenvironment is critical for initiating tumor-associated antigen-specific adaptive immunity and protection. Chemotherapeutic drugs have been extensively studied for their capacity to achieve protective immunity through ICD and DAMP release^32,33^. Mitoxantrone is an ICD inducer used in immunotherapies^34^. Cisplatin is not an ICD inducer but shows synergistic effects in immunotherapies^25^. In this study, we compare the ICD and DAMP release profiles of PAIIR to those of mitoxantrone and cisplatin in a breast cancer spheroid model. We further investigate the effects of DAMPs released from treated spheroids on the activation and pro-inflammatory phenotypes of DCs. Through these studies, we aim to emphasize the advantages of PAIIR over commonly utilized chemotherapeutics.

## 2 Results and Discussion

### 2.1 Spheroid formation of EMT6 cells

EMT6 is an immunogenic breast cancer cell line^35^. We used the EMT6 breast cancer cell line to generate the spheroids. While multiple techniques exist to form spheroids, such as ultralow attachment, hanging drops, magnetic levitation, and spinner flasks^36^, we employed the ultra-low attachment technique, a method we previously demonstrated for creating different-sized spheroids^37^. Drug resistance is often associated with the formation of a hypoxic core (necrotic core) ^38^. Therefore, we aimed to form large spheroids (>500 μm in diameter) since those larger than 500 μm in diameter exhibit gradient and zone formation ^39,40^. In this study, we cultivated and analyzed spheroid growth, utilizing them on day 7 post-formation when they reached approximately 800 mm in diameter **(Figure 2A and 2B)**. The surface area of the spheroids increased over time **(Figure 2C)** and displayed high circularity **(Figure 2D)** and solidity **(Figure 2E)** throughout the culture period.

**Figure 2.**
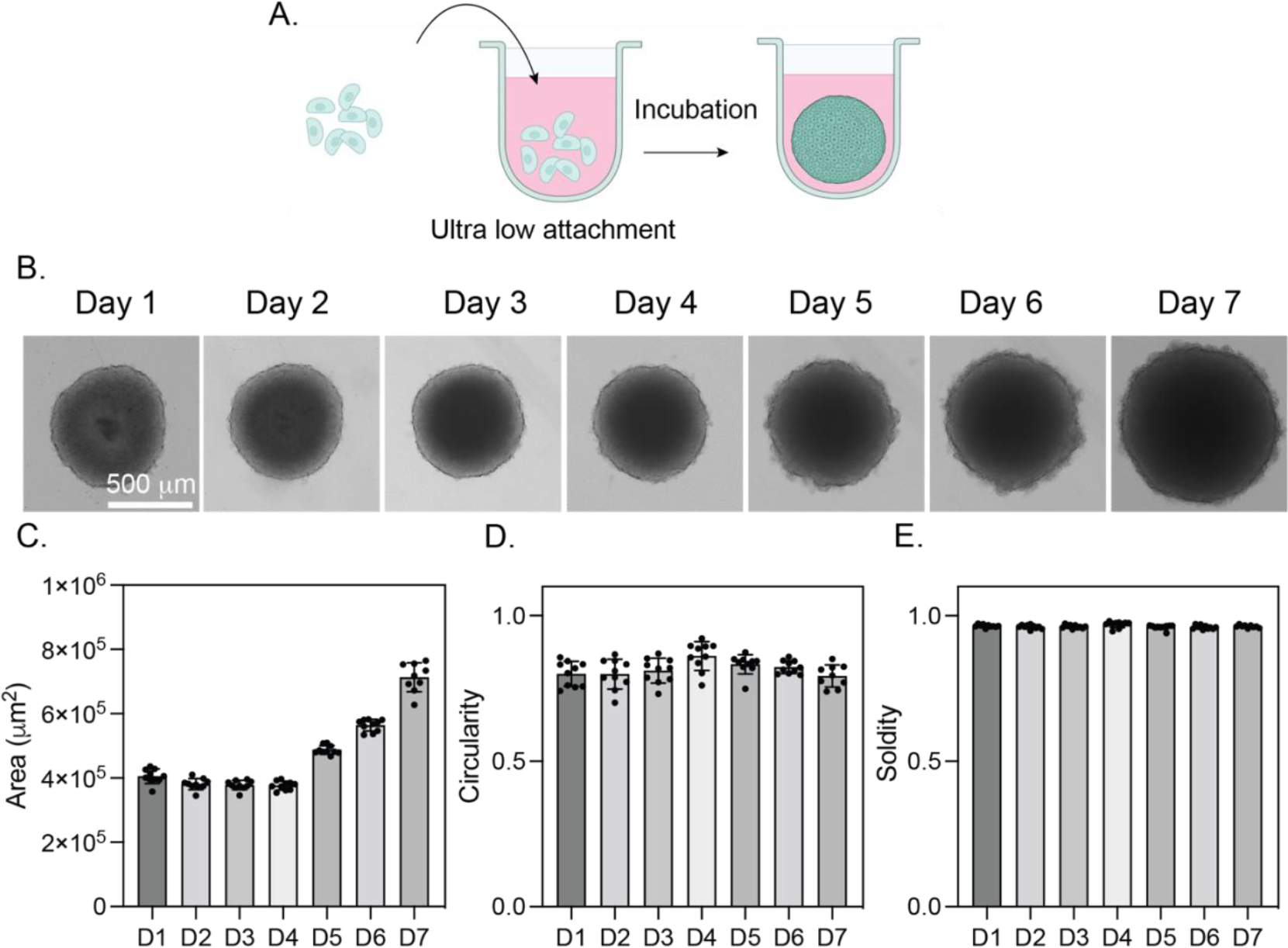
EMT6 spheroid formation and characterization. A. Schematic explanation of ultralow attachment spheroid formation. B. Representative brightfield images of EMT6 spheroids for 7 days. C. Spheroid area, D. Circularity, and E. Solidity of spheroids for 7 days. The scale bar is 500 μm.

### 2.2 Peptide-aggregation induces controllable cell membrane damage in EMT6 spheroids

We treated spheroids at day 7 for 24 hours using [II] concentrations of 0.5 mM, 0.75 mM, and 1 mM prepared in varying albumin concentrations (1%-0.5%-0.25%-0.125%-0.0625%). Across all [II] concentrations, an increased albumin content correlated with heightened cytotoxicity **(Figure 3A, 3B, and 3C)**. To determine if [II] triggered cell membrane damage within spheroids, we measured the release of lactate dehydrogenase (LDH)—a cytoplasmic enzyme released upon cell membrane damage ^41^. All [II] concentrations prompted LDH release **(Figure 3D, 3E, and 3F)**, indicating that aggregation kinetics directly affect the amount of cell death within the spheroid, not the type of cell death. However, increasing the peptide concentration from 0.5 mM to 0.75 and 1 mM did not induce significant cell death within the spheroids **(Figure 3A, 3B)**. This likely resulted from an elevated peptide-to-albumin ratio, which facilitates rapid assembly into non-cytotoxic fibrillar structures. The thioflavin T (ThT) assay is commonly utilized to detect β-sheet structures during Aβ fibrillization^42^. We observed the increasing intensity of ThT (represents increasing fibrilization) with 0.5 mM peptide prepared in 1% albumin. The ThT intensity and cell death profile were inversely correlated. As the peptide concentration rose, so did fibrilization levels, **(Figure 3G)**, indicating reduced cytotoxicity. Within a set concentration of albumin (1%), the peptides aggregate with albumin rather than forming an ordered, non-cytotoxic fibrillar state **(Figure 3H)**.

**Figure 3.**
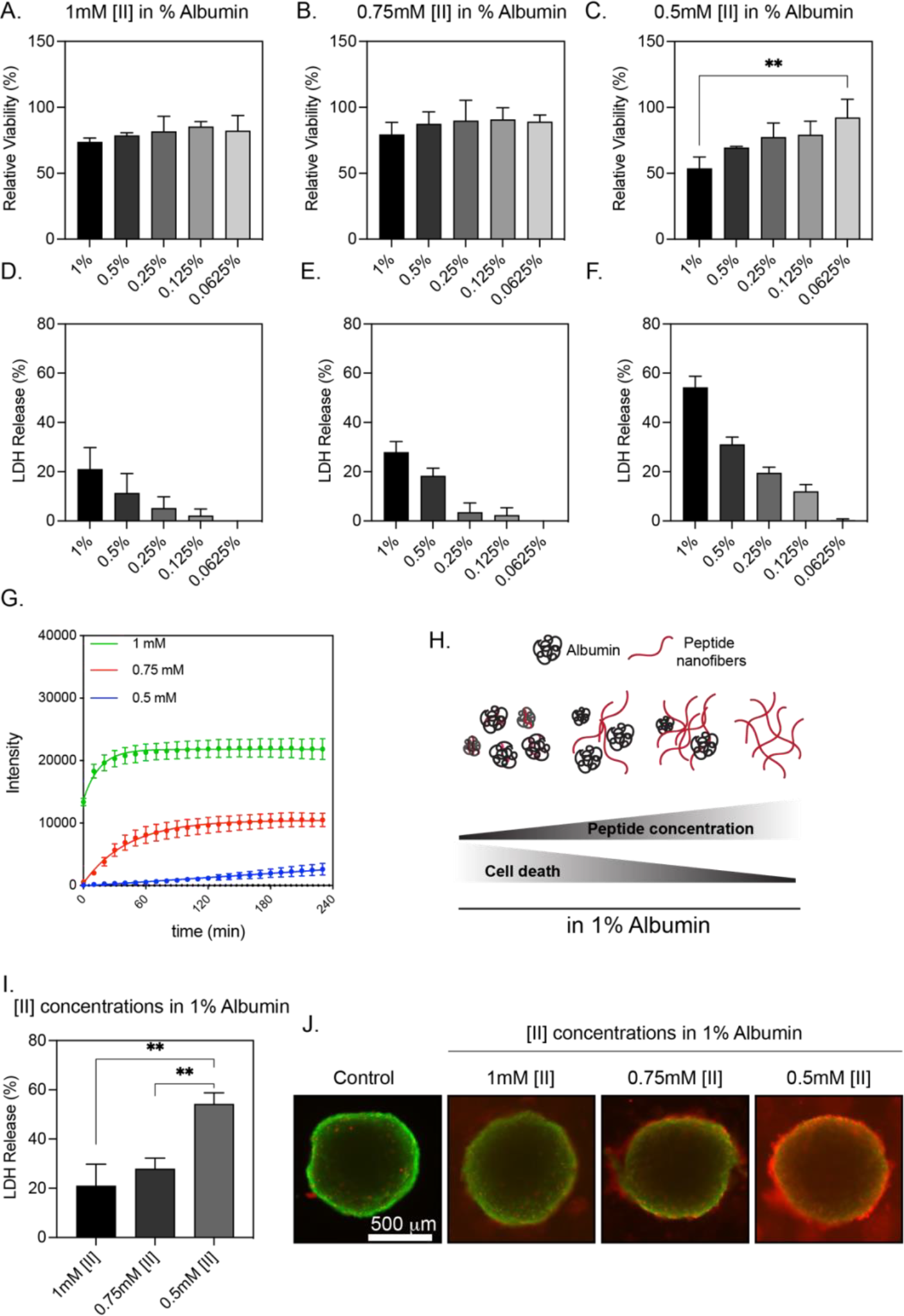
[II] Aggregation induced controllable membrane damage in the EMT6 spheroid model. Relative viability measurements of A. 1 mM, B. 0.75 mM, and C. 0.5 mM [II] peptide were prepared in different albumin concentrations at 24h. LDH release measurements of D. 1 mM, E. 0.75 mM, and F. 0.5 mM [II] peptide were prepared in different albumin concentrations at 24h. G. ThT measurement of 1 mM, 0.75 mM, and 0.5 mM [II] peptide prepared in 1% albumin. H. Schematic showing the transition from the aggregated state to the fibrillar structure with increasing peptide concentration. I. Comparison of LDH release of 1 mM, 0.75 mM, and 0.5 mM [II] peptide prepared in 1% albumin. J. Representative calcein (live cells, green) and propidium iodide (membrane damaged cells, red) images of spheroids treated with 1 mM, 0.75 mM, and 0.5 mM [II] peptide prepared in 1% albumin. Statistical analysis was performed with a one-way ANOVA test; data are presented as mean ± S.D., **p<0.01. The scale bar is 500 μm.

Among the groups studied, 0.5 mM [II] peptide in 1% albumin induced significant (**p<0.01) cell death within the spheroids at 24h **(Figure 3C)**. This result also showed the controllability of cell death within the spheroids by modulating the aggregation of the peptide with albumin. LDH release was highest with 0.5 mM [II] prepared in 1% albumin **(Figure 3I)**, in line with the viability experiments.

Subsequent comparisons of LDH release from spheroids treated with [II] peptides (1 mM, 0.75 mM, and 0.5 mM) in 1% albumin demonstrated that 0.5 mM [II] peptide induced significantly (**p<0.01) higher LDH release compared to other conditions **(Figure 3J)**. Findings were confirmed with imaging with a fluorescent live/dead assay. [II] peptide-treated spheroids were stained with calcein (live cells, green) and propidium iodide (membrane-damaged cells, red). Spheroids treated with 0.5 mM [II] in 1% albumin had the highest propidium iodide uptake, indicating membrane damage consistent with the LDH and viability data. These results underscore that [II] can induce tunable membrane damage in EMT6 spheroids via albumin control.

### 2.3 EMT6 spheroids have higher chemotherapeutics resistance than cell culture monolayers

Antigens spread from dying cells can initiate adaptive immunity, specifically when accompanied by DAMPs that provide inflammatory and immunogenic signals which determine the efficacy of the activation of innate immunity.^43^ To compare the efficacy of released signals from cells dying by [II] peptides and chemotherapeutics, we studied the 50% cell death ratio within spheroids (IC_50_ values) in 24h with all groups.

Both monolayer and spheroid cultures of EMT6 cells were treated with varying concentrations of mitoxantrone and cisplatin to determine their respective IC50 values.. Mitoxantrone had an IC_50_ value of 1.382 μM for monolayer culture and 47.31 μM for spheroid culture **(Figure 4A and 4B)**. In comparison, cisplatin had 14.32 μM for monolayer culture and 28.49 μM for spheroid culture **(Figure 4A and 4B)**. Resistance of cancer spheroids to chemotherapeutics has been shown by us^37,44^ and others^45–47^ before. Our results also show that EMT6 spheroids are resistant to mitoxantrone and cisplatin. These results show that spheroid models exhibit drug resistance similar to that in in vivo tumors, and offer a high-throughput platform for studying new therapeutics.

**Figure 4.**
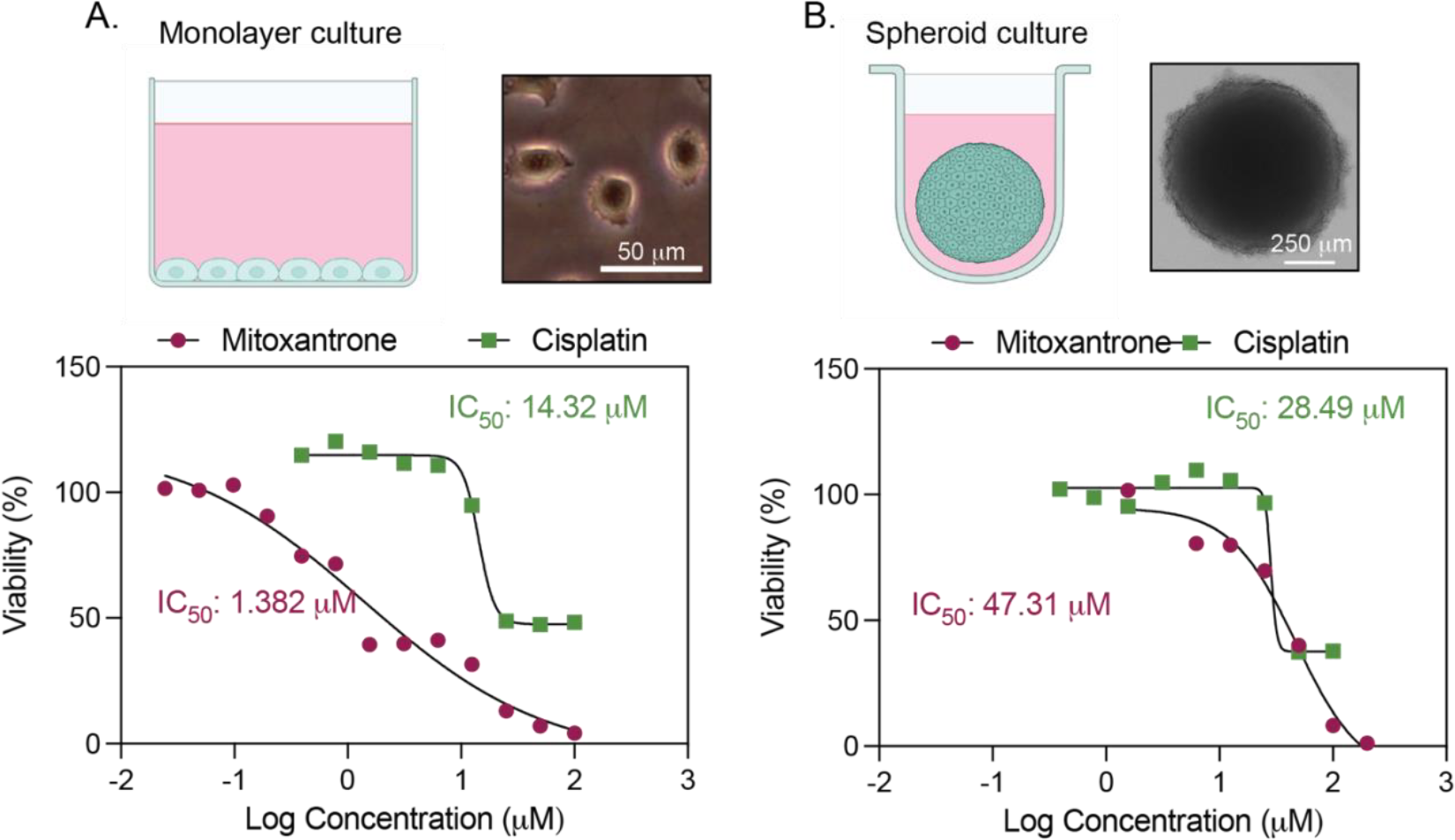
IC_50_ values of chemotherapeutics in monolayer and spheroid cultures. A. IC_50_ values of mitoxantrone and cisplatin in monolayer culture at 24h. The scale bar is 50 μm. B. IC_50_ values of mitoxantrone and cisplatin in spheroid culture at 24h. The scale bar is 250 μm.

Despite their potency, traditional chemotherapeutics have severe side effects when delivered systemically^48^.One of the most significant limitations is the suboptimal drug accumulation in solid tumors. Recent efforts have focused on the local delivery of chemotherapeutics over than systemic administration^48^. However, within the tumor microenvironment (TME), multiple factors contribute to drug resistance, including the upregulation of anti-apoptotic genes^49^. In contrast, the peptide aggregation model we engineered does not target a specific receptor to initiate ICD and DAMP release; therefore, it presents a unique model to overcome acquired resistance against chemotherapeutics.

### 2.4 PAIIR induced DAMP release is superior to chemotherapeutics

DAMPs have been proven necessary for the immunogenicity of dying cells^33,50,51^. We previously showed [II] aggregation-induced ICD and the release of DAMPs in different cell lines^18^. To identify differences in DAMP release, we treated EMT6 spheroids on day 7 for 24h with 0.5 mM [II] peptide in 1% albumin, mitoxantrone 50 μM, and cisplatin 30 μM (near their 50% spheroid death concentrations for all conditions) and compared the DAMP release profiles. LDH release was significantly (****p<0.0001) higher in [II] peptide- and mitoxantrone-treated spheroids than in cisplatin-treated spheroids **(Figure 5A)**, indicating higher membrane damage with peptide or mitoxantrone. The spheroid supernatant was analyzed 24 hours post treatment for the release of DAMPs. A diverse set of molecules are released upon ICD, such as extracellular ATP, nucleic acids, and high mobility group box 1 (HMGB-1)^50,52^. Extracellular release of ATP acts on purinergic receptors^8^ on DCs to secrete IL-1β and activate tumor antigen-specific CD8+ T cells, an effect that is abolished in the absence of purinergic receptors^53^. We detected significantly (****p<0.0001) higher extracellular ATP levels upon [II] treatment compared to chemotherapeutics **(Figure 5B)**. Although past studies have shown detectible extracellular ATP with lower concentration mitoxantrone treatments^34,54^, neither mitoxantrone nor cisplatin yielded observable ATP in our study^15^ **(Figure 5B)**. The concentration of chemotherapeutics changes their ability to induce ICD and DAMP release^15^; therefore, the lack of extracellular ATP could be due to the concentration used in our study.

**Figure 5.**
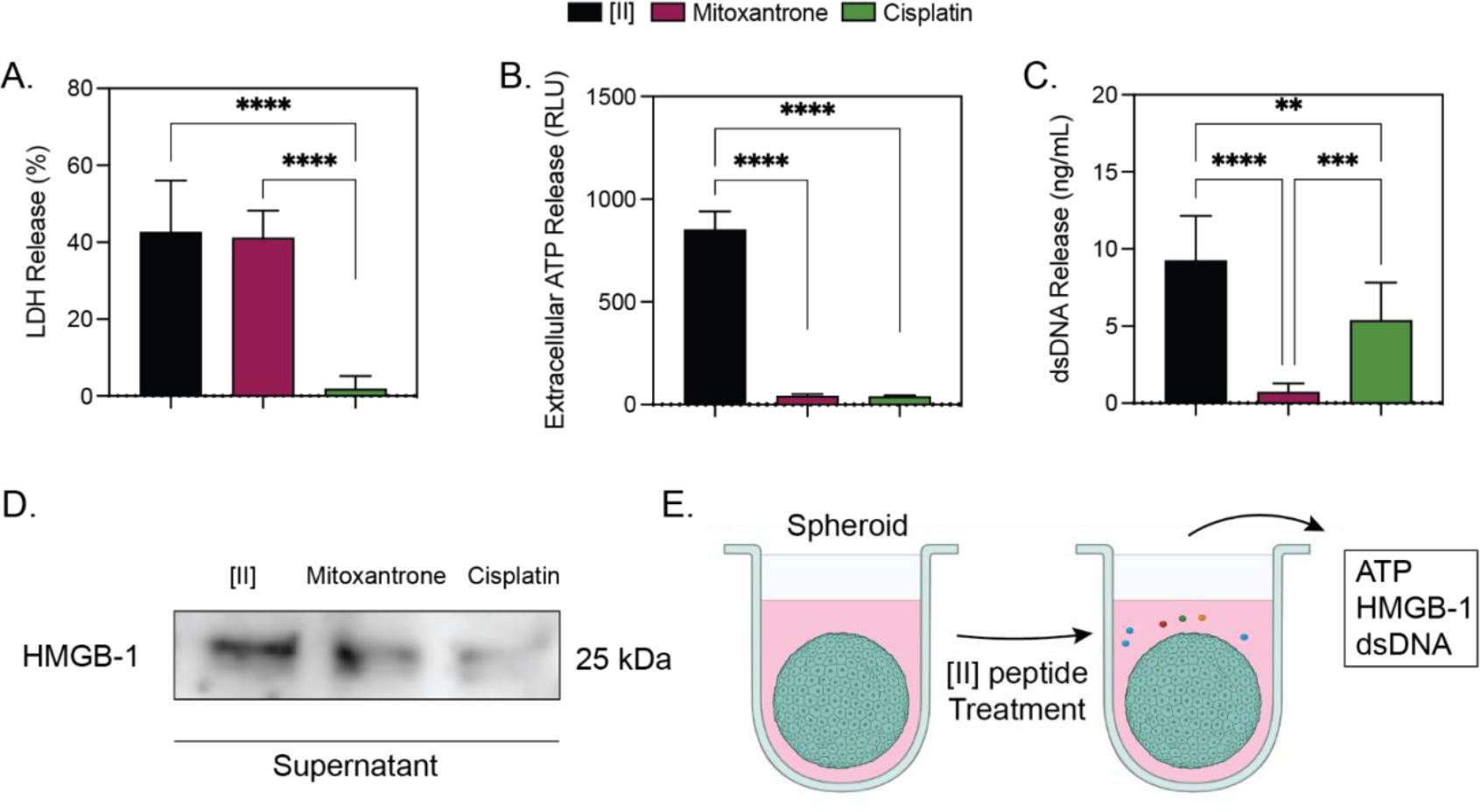
DAMP release from EMT6 spheroids. A. Release of LDH, B. Extracellular ATP, C. dsDNA, and D. HMGB-1 after [II], mitoxantrone, and cisplatin treatment at 24h. E. Schematic represents the DAMPs released by [II] aggregation. Statistical analysis was performed with a one-way ANOVA test; data are mean ± S.D., **p<0.01, ***p<0.001, ****p<0.0001.

Pattern recognition receptors (PRRs), such as toll-like receptors (TLRs) on immune cells, are specialized to detect different “patterns,” including DAMPs^55^. For example, nucleic acid sensing is critical for inducing an antiviral immune response against DNA and RNA viruses^56^. Nucleic acids initiate a robust type I interferon response, which is crucial for antitumor immunity^57^. Specifically, double-stranded DNA (dsDNA) increases the expression of genes in DCs responsible for antigen processing and presentation^58^. We detected significantly higher dsDNA release upon [II] treatment compared to chemotherapeutics **(Figure 5C)**. Cisplatin also induces dsDNA release, possibly due to its role in inducing double-strand DNA breaks^59^. HMGB-1 is a nuclear protein that is released during ICD. Extracellularly, it can engage TLR4 and receptor for advanced glycation end products (RAGE) and stimulate pro-inflammatory cytokine production^60^. We detected higher levels of HMGB-1 in the extracellular matrix of spheroids treated with [II] than in those treated with chemotherapeutics **(Figure 5D)**. Summarily, [II] aggregation induced the release of ATP, dsDNA, and HMGB-1 **(Figure 5E)** and was superior to chemotherapeutic agents in both variety and quantity.

The effects of individual DAMPs on immune cells and their receptor interactions have been extensively studied. However, recent studies have highlighted the synergistic effects of DAMPs on the activation of immune cells. For example, silencing HMGB-1 only partially reduced the protective capacity of oxaliplatin-induced ICD in a prophylactic vaccination model^33^. The release of ATP and HMGB-1 in the early stage of ferroptosis was protective, whereas depletion of ATP at the late stage abolished this protection in a prophylactic vaccination model^51^. Given the synergistic effects of DAMPs, the [II] aggregation-induced DAMP release profile is promising for activating innate immune cells such as DCs.

### 2.5 [II] peptide aggregation-induced DAMP release activates BMDCs

Proper activation of DCs is crucial for the initiation of antigen-specific adaptive immunity. Several DAMPs and their effects on activating DCs have been studied: ATP and HMGB-1 for DC recruitment and activation^52^ and nucleic acids for antigen processing and presentation^58^. Bone marrow differentiated DCs (BMDCs) were utilized to analyze the effect of released DAMPs on DC cytokine release. We treated the spheroids with [II] peptide, mitoxantrone, and cisplatin for 24h at the given concentrations, then transferred their supernatants (spheroid-conditioned medium) onto BMDCs for analysis of cytokine release profiles at both 6 and 24h **(Figure 6A)**.

**Figure 6.**
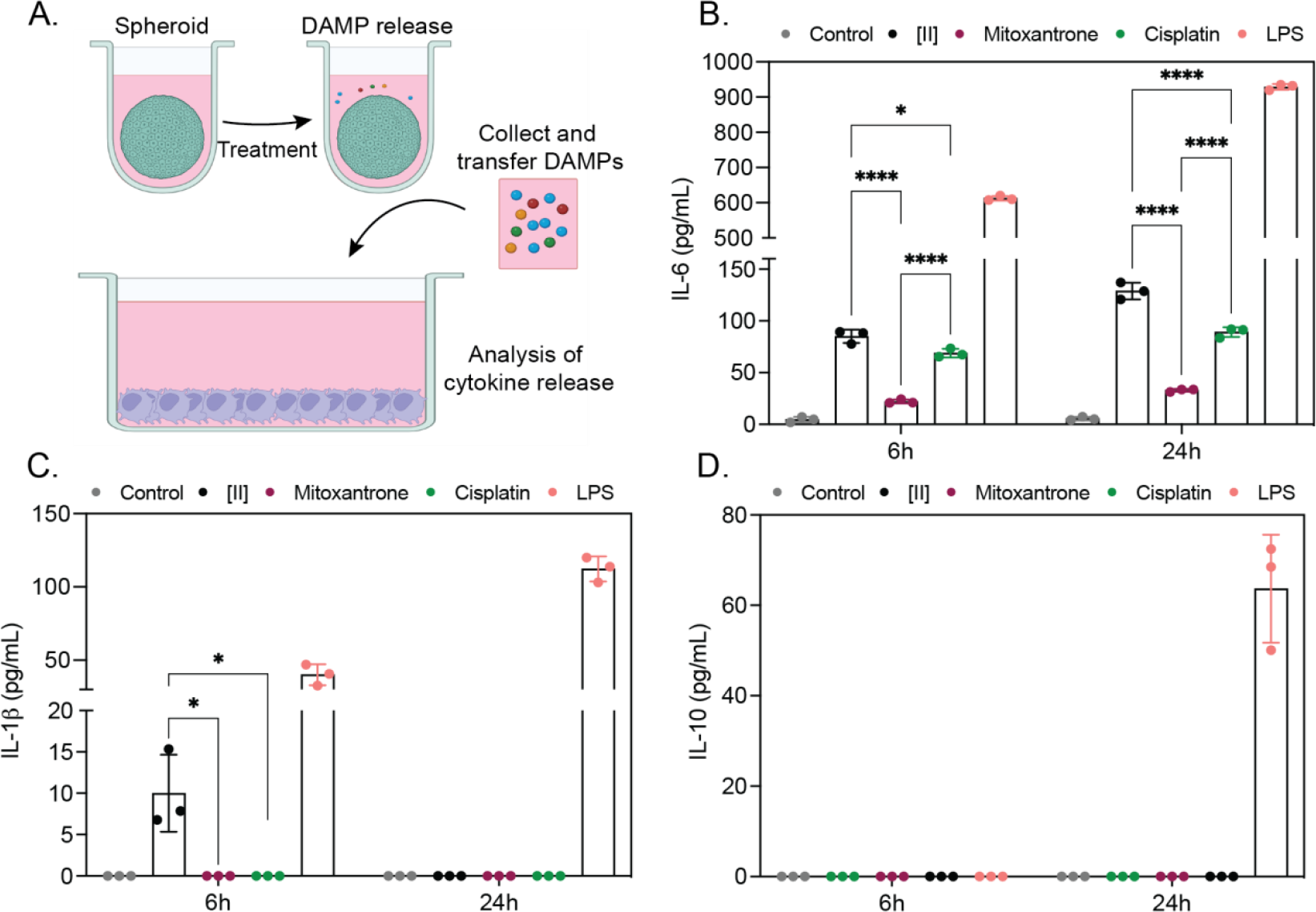
[II] Aggregation-induced DAMP release activates BMDCs. A. Experimental plan for the treatment of BMDCs with tumor spheroid conditioned mediums. B. Release of IL-6, C. IL-1β, and D. IL-10, from BMDCs at 6h and 24h after tumor spheroid conditioned medium treatment. Statistical analysis was performed with a two-way ANOVA test; data are mean ± S.D., *p<0.1, ****p<0.0001.

We identified a significant (*p<0.1, ****p<0.0001) increase in pro-inflammatory cytokine IL-6 secretion when BMDCs were cultured with [II]-treated spheroid conditioned medium **(Figure 6B)**. IL-1β was only released from BMDCs cultured with [II] treated spheroid-conditioned medium **(Figure 6C)**. This release is attributed to the presence of extracellular ATP from [II] treatment **(Figure 5B)**, as previous studies have shown that extracellular ATP induces IL-1β release from DCs ^53^. However, the degradation of ATP extracellularly^61^accounts for the absence of IL-1β release at 24h **(Figure 6C)**. We detected no anti-inflammatory cytokine IL-10, under any of the conditions.. Lipopolysaccharide (LPS) was used as a positive control for BMDC activation^62^.

### 2.6 [II]-peptide treated spheroid conditioned medium is non-toxic to BMDCs

Although Mitoxantrone is a recognized ICD inducer^25^, we did not detect its effect on DAMPs activating BMDCs **(Figure 6B and 6C)**. To further understand this, we measured the viability of BMDCs after 24h exposure to spheroid-conditioned mediums **(Figure 7A)**. Among the groups, only mitoxantrone induced additional significant cytotoxicity in BMDCs **(Figure 7B)**, explaining the low levels of cytokine release from BMDCs **(Figure 6B and 6C)**. This effect was due to the high mitoxantrone concentrations required to induce 50% cytotoxicity **(Figure 4B)** in the spheroid model.

**Figure 7.**
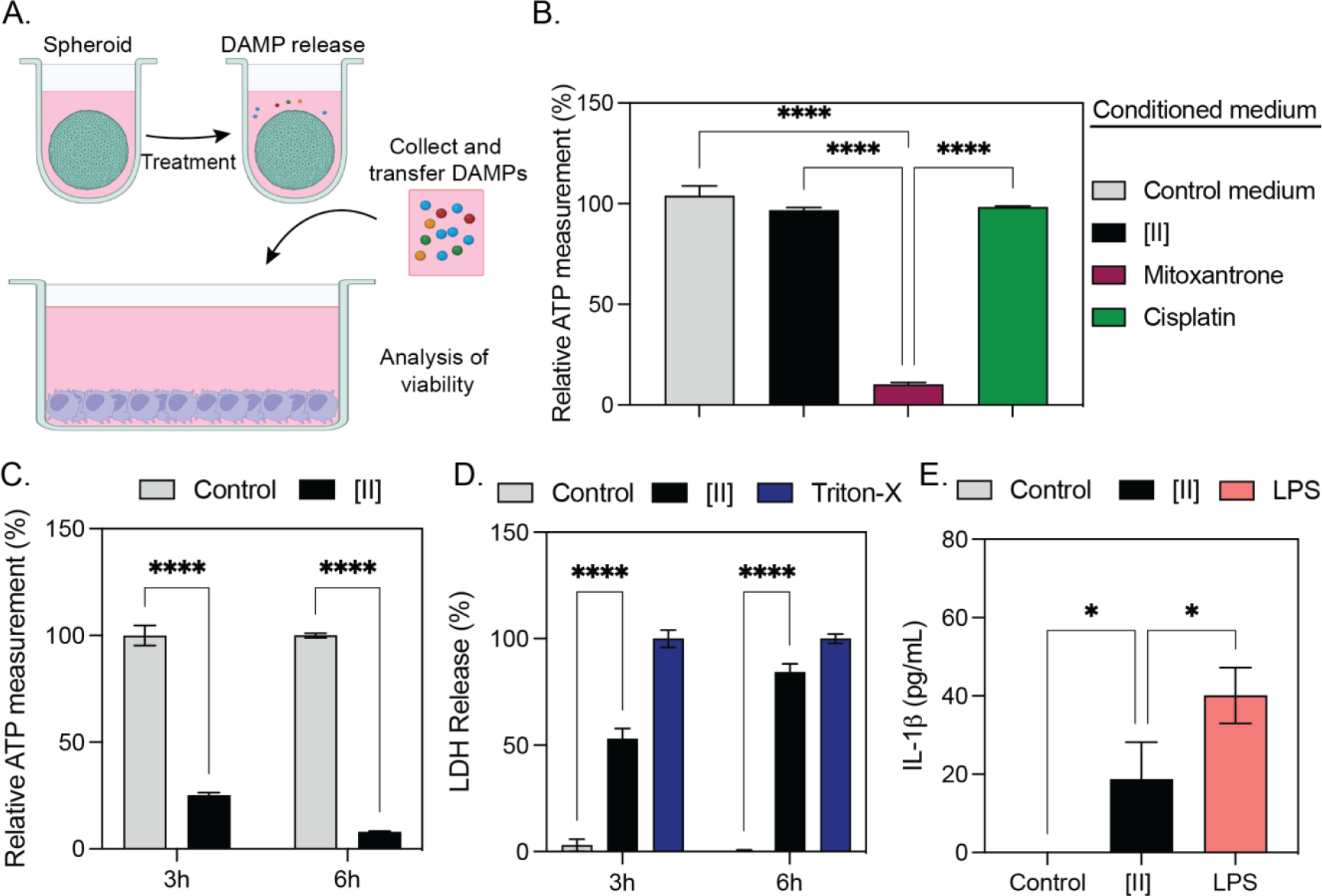
[II]-treated spheroid conditioned medium is not cytotoxic to BMDCs. A. Schematic of the design for the viability measurement of BMDCs. B. Viability measurement of BMDCs after incubation with spheroid-conditioned mediums at 24h. C. Viability and D. LDH and E. IL-1β release (6h) of BMDCs after [II] peptide treatment. Statistical analysis was performed with one-way ANOVA (B and E) and two-way ANOVA tests (C and D); data are mean ± S.D., ****p<0.0001.

As stated previously, chemotherapeutics have side effects and are toxic to normal tissues when administered systemically^63^. While local/intratumoral delivery of chemotherapeutics reduces the side effects, the severity of side effects depends on required concentrations and diffusion parameters^64^. Conditioned medium from [II] peptide had no cytotoxicity to BMDCs, showing that [II] effect would be contained locally **(Figure 7B)**, as we have shown before^18^. In the case of an intratumoral treatment strategy, the initial injection will also affect multiple locally present cell types, including DCs. To understand the effects of [II] on DCs we treated BMDCs with [II] peptide. We found that [II] aggregation induces time-dependent cell death **(Figure 7C)** and cell membrane damage **(Figure 7D) and** prompts IL-1β secretion from BMDCs **(Figure 7E)**. IL-1 family members such as IL-1β have been discussed as canonical DAMPs and inducible DAMPs capable of activating immune cells^43,65^. Therefore, in addition to DAMPs such as ATP, dsDNA, and HMGB-1, [II] peptide also releases inducible DAMPs.

### 2.7 [II] peptide aggregation increases antitumor cytokines in breast cancer mouse model

To test the efficacy of [II] peptide aggregation in vivo, we formed a breast tumor model using a murine EMT6 cell line. C57BL/6 mice were inoculated with EMT6 cells to form palpable tumors by day 8. Results **(Figure 3)** indicate that there is an increasing trend of ICD with increased albumin concentrations at a concentration of 0.5 mM [II]. Thus, to observe the effects more clearly in vivo, we conducted the study using 0.2% and 2% albumin concentrations, allowing us also to monitor potential systemic toxicity from these distinct concentrations.

The selected [II] combinations were administered intratumorally every two days for a duration of 14 days **(Figure 8A)**. On day 22, the mice were euthanized for analysis of the cytokine profiles in the tumor microenvironment. As expected, the delayed aggregation of [II] in 2% MSA showed more potent effects compared to the control group and the group treated with 0.2% MSA, evidenced by significantly lower tumor volume at day 22 (*p<0.1). Tumors treated with 0.5 mM [II] in 0.2% and control had no statistically significant difference **(Figure 8B)**. Additionally, the analyzed levels of IL-1β (**p<0.01) and IFN-*γ* (*p<0.1) levels were significantly increased upon [II] peptide treatment with 2% MSA **(Figure 8C)** compared to control and 0.2% MSA conditions. However, IL-6 was significantly elevated in the control group **(Figure 8C)**. No significant differences were detected for IL-10 and IL-12p70 among the samples.

**Figure 8.**
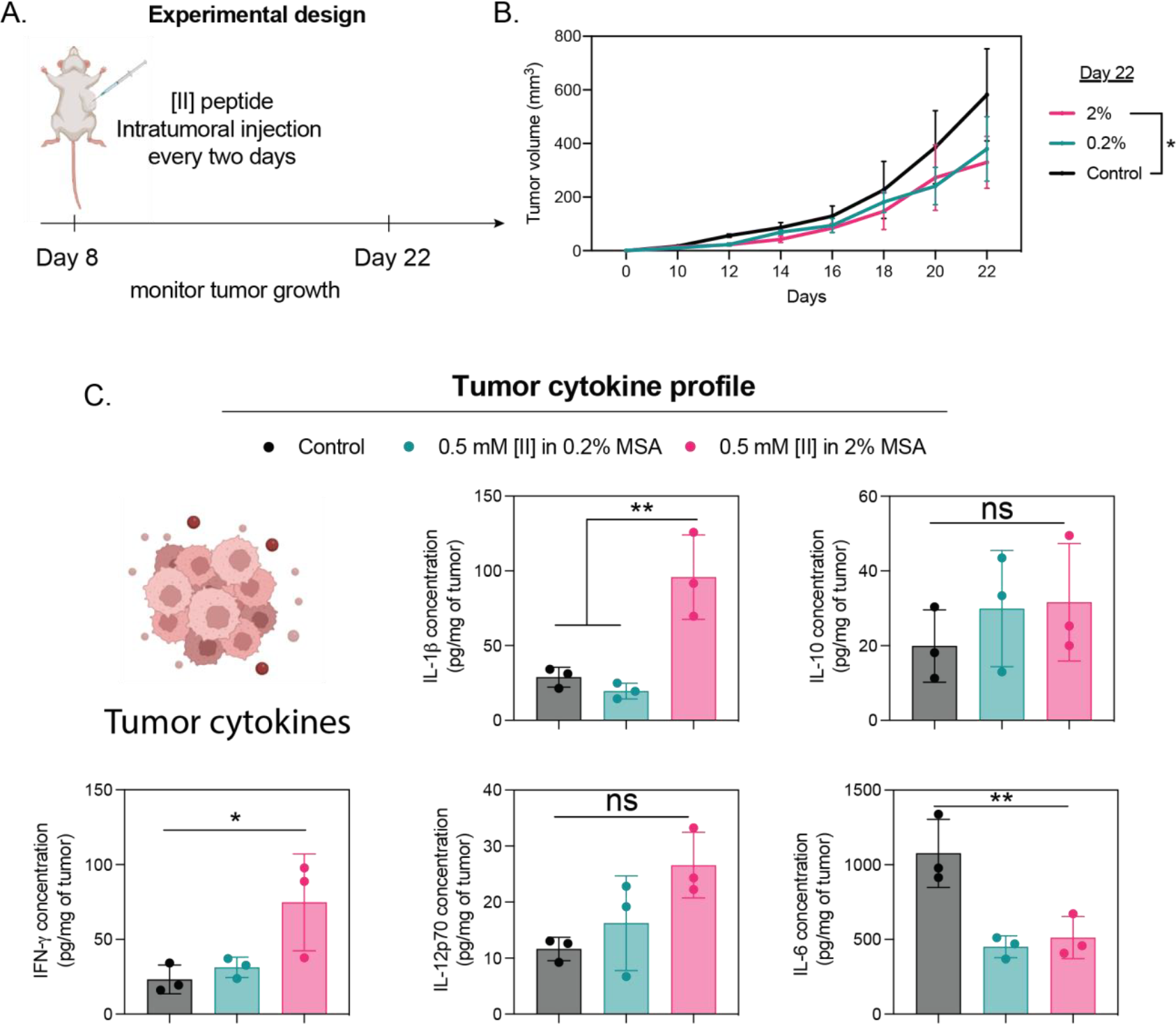
[II] peptide aggregation increases antitumor cytokines in breast cancer mouse model. A. Experimental design of tumor formation and treatment strategy. B. Tumor volume measurement for 22 days. C. Tumor microenvironment cytokine analysis at day 22. Statistical analysis was done with one-way ANOVA (B and E) and two-way ANOVA tests (C and D); data are mean ± S.D., *p<0.1, **p<0.01.

The cytokines IL-1β and IFN-*γ* can trigger inflammatory responses and stimulate activation of immune cells, such as T cells and natural killer (N.K.) cells^66–68^. The release of pro-inflammatory cytokines, including TNF-α, IFN-γ, and IL-1β, promotes the activation of myeloid dendritic cells and the differentiation of T cell subsets^69^. In our results, the identification of significantly higher IL-1b and levels with intratumoral injection of 0.5 mM [II] with 2% MSA (Figure 8C) indicates higher activation of antitumor immune response in vivo.

Prior research has demonstrated the detrimental effects of IL-10 on tumor progression, as it hinders antigen-presenting cell function by suppressing MHC and costimulatory molecules, thereby inducing immune suppression or tolerance^70,71^ Conversely, IL-12p70 has been found to enhance the production of interferon-gamma, a potent cytokine with antitumor properties^72^. The heightened levels of IL-6 have been associated with metastasis promotion and inhibition of antitumor immune responses^73,74^

In our study, the notable decrease in IL-6 levels treated with [II] could potentially augment the targeted immune response against the tumor. This suggests that a combination of 0.5 mM [II] with 2% MSA may facilitate a stronger immune response within the microenvironment of the tumor, as indicated by cytokine profiling outcomes.

### 2.8 Histological investigation of toxicity in liver and kidney

Drug-induced kidney and liver injuries are the most common side effect that has been observed in patients undergoing cancer immunotherapies, relatively early during the treatment in the first 30-120 days.^75,76^ The severity of the side effects can cause the discontinuation of immunotherapies in some cases.^77–79^Identification and assessment of histological damage in immunotherapy-associated liver and kidney injuries has been valuable in both differential diagnosis and determining the severity of the damage.

At the end of the treatments, mouse kidneys were for histological investigation. Although the mice received a total of seven injections of the [II] peptides intratumorally, no significant damage was observed in either of the organs (**Figure 9**).

**Figure 9.**
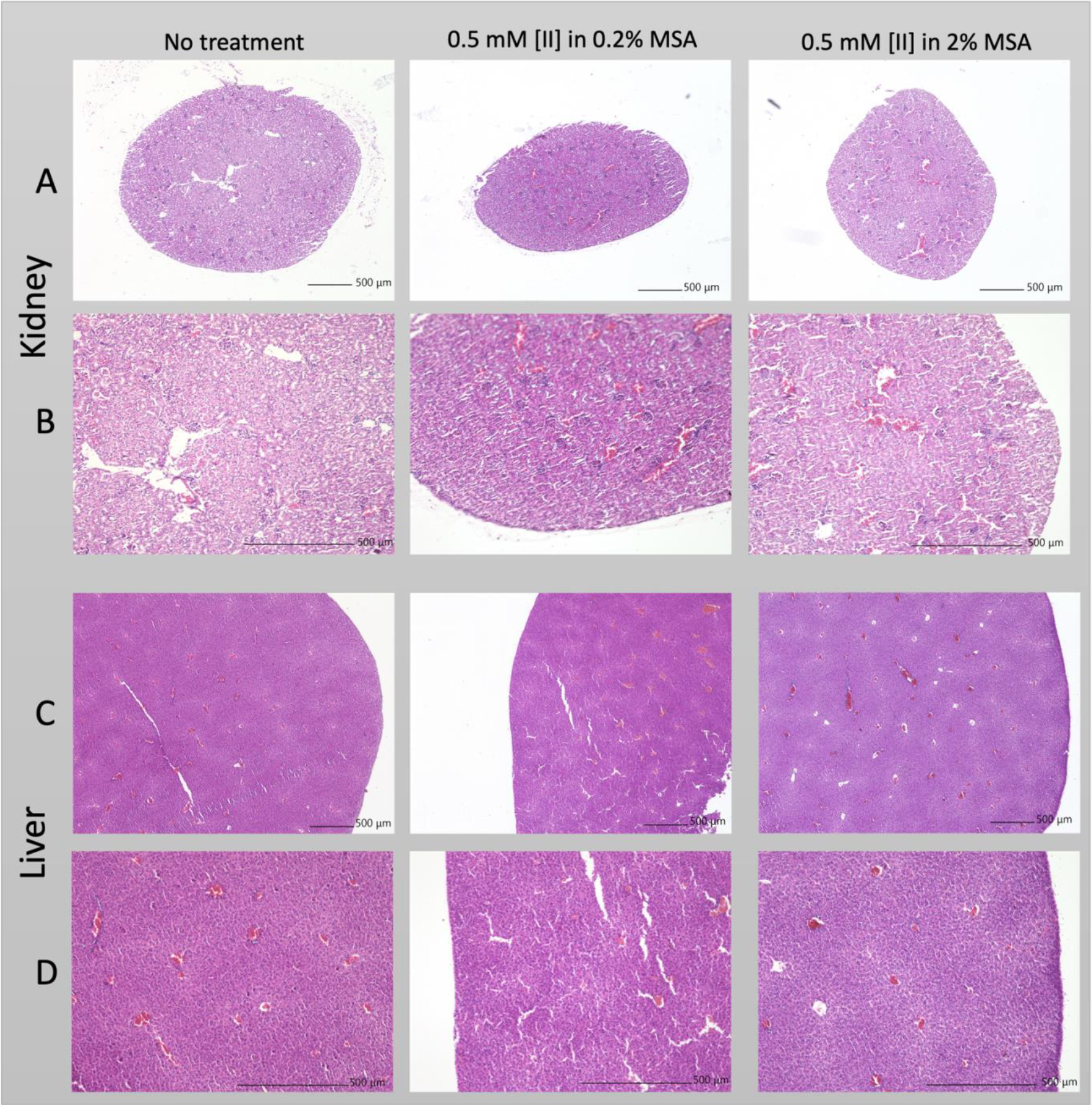
Histology of [II]-treated mice kidney and liver. H&E stained histology of A. kidney in 4X and B. 10X magnifications, C. liver in 4X and D. 10X magnifications. All scale bars = 500 μm.

Although immune-mediated processes may trigger auto immune diseases, renal side effects of immunotherapy are relatively infrequent^80^. The thorough studies comparing clinicopathologic features between immunotherapy-induced liver injury and classic autoimmune liver diseases identifiy some similarities, such as predominantly lobular hepatitis with milder portal inflammation, but there are also obvious differences between the two conditions^76,81^. Similarly, histologic observation of kidney damage in patients undergoing immunotherapy has revealed acute tubulointerstitial nephritis as the most common manifestation.^82,83^ regardless of the extent of glomerulosclerosis severity^84^. Observation of no significant damage in the histological images of the organs from the treated mice indicates that [II] did not show toxicity on these organs in this treatment. More extensive studies, including the enzymes related to kidney and liver health, should be investigated for more accurate information.

## 3 Conclusions

We used a breast cancer spheroid model to examine the effects of peptide aggregation-induced ICD alongside recognized chemotherapeutics. Our results showed that [II] aggregation-induced ICD can be controlled and is superior to chemotherapeutics in terms of the type and amounts of DAMPs. DAMPs released upon [II] treatment activated DCs to secrete pro-inflammatory cytokines that have a role in orchestrating antitumor immunity. Furthermore, the transfer of [II] peptide solution did not induce further cytotoxicity, highlighting that [II] aggregation-induced ICD and DAMP release would be contained locally.

Considering that different chemotherapeutic concentrations can alter the immunogenicity and DAMP release of dying cells^15,33,85^, it is necessary to use physiologically relevant models to study ICD and DAMP release. Classical monolayer culture often falls short in evaluating the efficacy of new treatment strategies for in vivo applications and their progression to clinical trials. Spheroids offer physiological relevance to solid tumors and provide a high-throughput system to test various conditions to select promising candidates for further studies. Their resistance to chemotherapeutics impacts cell death dynamics, making it essential to use models that closely resemble physiological conditions. Our study revealed the peptide aggregation model’s superiority in DAMP release and BMDC activation within the breast cancer spheroid model. Relying solely on monolayer cultures might not yield accurate physiological results. Hence, spheroid models present a more appropriate framework to evaluate treatment strategies and test new therapeutic interventions.

## 4 Materials and Methods

### Statistical analysis

Data were analyzed by using GraphPad Prism Version 9.3.1. Adobe Illustrator 2021 was used for figure preparation. Figure 3A, 3B, 3C, 7B, and 7C were normalized to the mean of untreated control. Figure 3D, 3E, 3F, 3I, 5A and 7D were normalized to the mean of Triton-X treated control group for LDH release (Maximum LDH release). At least three biological replicates were used. All data are presented as mean ± standard deviation (S.D.). Statistical analysis was done by one-way analysis of variance (ANOVA) and two-way ANOVA with Tukey’s multiple-comparison test. **** p<0.0001, *** p<0.001, ** p<0.01, * p<0.1.

### Materials

9-fluorenylmethoxycarbonyl (Fmoc) protected amino acids, [4-[α-(2’,4’-dimethoxyphenyl) Fmoc aminomethyl] phenoxy] acetamidonorleucyl-MBHA resin (Rink amide MBHA resin), Oxyma, N, N’-Diisopropylcarbodiimide (DIC), piperidine, Trifluoroacetic acid (TFA), dimethylformamide (DMF), dichloromethane (DCM) were purchased from Gyros Protein Technologies. Triisopropylsilane, Acetic anhydride, Congo Red dye, and pyrene were purchased from Sigma-Aldrich. Deionized water (resistance of 18.2 MΩ.cm) was used during the experiments. Peptides were purchased from Biomatik (Cambridge, Ontario, Canada).

### Thioflavin T (ThT) Assay

ThT (Sigma-Aldrich) was used to understand the aggregation kinetics of [II] peptide. ThT measurement included 1% albumin for 0.5 mM, 0.75 mM, and 1 mM [II] peptide. At first, fresh albumin (40 mg/mL, 4 %w) was dissolved in 1x PBS and diluted into 1% in 1x PBS. Then, stock peptides (10 mM in water) were diluted (0.5 mM) individually. For ThT measurements, 5 μL from 400 μM ThT in PBS was added to 195 μL of 0.5 mM KFFIIK and EFFIIE individually. Then, each peptide was mixed and read with BioTek Neo2SM microplate reader for 4 h with 10 min intervals (Ex: 440, Em:480 with gain:90).

### Cell culture and reagents

Mouse breast cancer cell line EMT6 was cultured in a humidified incubator at 37°C supplied with 5% CO2. EMT6 cells were cultured in Roswell Park Memorial Institute (RPMI) media (SIGMA R8758) supplemented with 10% FBS (Hyclone SH30910.03), 6 mM HEPES, 50 μM 2-mercaptoethanol and 1% antibiotics; penicillin (100 U/mL), and streptomycin (100 μg/mL) (Thermo Fisher 15240062). T75 flask (TPP 90076) was used to culture cells, and cells were passaged upon 85% confluency using trypsin (Sigma 59418C). The media was changed every 2 days.

### Spheroid formation and characterization

EMT6 spheroids were formed through ultra-low attachment technique in U-bottom 96 well plates (CELLTREAT 229590). Briefly, wells of the U-bottom 96 well plate were treated with an anti-adherence rinsing solution (STEMCELL Technologies 07010) and centrifuged at 1300g for 5 min. Then, the solution was aspirated, and surfaces were washed with the basal medium. Then, wells were seeded with 20.000 EMT6 cells, and plates were centrifuged at 100g for 3 min. Media was changed every day by carefully replacing 100 μL with fresh medium. Spheroids were imaged every 24h and later analyzed for their area, diameter, circularity, and solidity by using ImageJ.

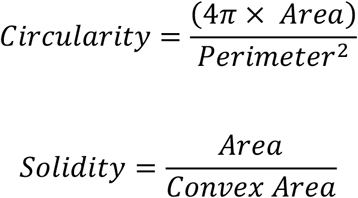

### Peptide and chemotherapeutics preparation and treatment

EFFIIE and KFFIIK were prepared in indicated concentrations in different albumin concentrations. We previously established the preparation technique of [II] peptide^18^. Briefly, individual charged counterparts were prepared in albumin solutions and mixed for 30 minutes before treatment. Mitoxantrone (Millipore Sigma M6545) and cisplatin (Millipore Sigma 232120) were used at indicated concentrations and prepared in normal cell culture medium.

### Lactate dehydrogenase (LDH) measurement

Cytoscan-LDH cytotoxicity assay kit (G-BIOSCIENCES 786-210) was used to measure LDH release from treated spheroids. Briefly, at the end of the 24h treatments, supernatants were collected after treatments and mixed with LDH reaction mixture for 20 min at 37°C. The reaction was stopped using the stop solution, and absorbance was measured at 490 nm and 680 nm. Triton-X was used to lyse the spheroids and used as a positive control for LDH release. 680nm background signal was subtracted, and relative LDH release was calculated based on the positive maximum LDH release control group.

### Extracellular ATP measurement

CellTiter-Glo 2.0 solution (Promega G9248) was used to measure the extracellular ATP released from spheroids. Briefly, at the end of 24h treatment, supernatants were mixed with CellTiter-Glo 2.0 solution and incubated for 5 min at room temperature. Then, the luminescent signal was measured with BioTek Neo2SM microplate reader according with the manufacturer’s instructions.

### Extracellular dsDNA measurement

Extracellular dsDNA release from spheroids was measured using AccuGreen™ High Sensitivity dsDNA Quantitation Kit (BIOTIUM 31066-T). Briefly, at the end of 24h treatment, supernatants from spheroids were collected, and the released amount of dsDNA was measured accordingly with the manufacturer’s instructions. A standard curve was generated and used to quantify the released dsDNA amounts.

### Immunoblotting reagents

Acrylamide/Bis-acrylamide, 30% solution (Sigma A3699), 1.5 M Tris-HCl, pH 8.8 (Teknova T1588), Tris HCl Buffer 0.5 M solution, sterile pH 6.8 (Bio Basic SD8122), Ammonium persulfate (Sigma A3678), UltraPure 10% SDS (Invitrogen 15553-027), TEMED (Thermo Fisher Scientific 17919), Dithiothreitol (DTT) (BIO-RAD 1610610) Tris Base (Fisher Bioreagents BP152), Glycine (Fisher Bioreagents BP381), 4x Laemmli sample buffer (BIO-RAD 1610747), TWEEN 20 (Sigma P9416), Mini Trans-Blot filter paper (BIO-RAD 1703932), Nitrocellulose Membranes 0.45 μm (BIO-RAD 1620115), Western Blotting Luminol Reagent (Santa Cruz sc-2048).

### Procedure

Protein samples were diluted in Laemmli buffer and boiled for 5 min at 96 °C. Proteins were then separated by sodium dodecyl sulfate-polyacrylamide gel electrophoresis (SDS-PAGE) gels (15%) and transferred to nitrocellulose membranes. After transfer, the membranes were blocked in 5% milk in Tris Buffered Saline with 0.1% Tween 20 (TBST). Blots were then incubated with HMGB-1-HRP (BioLegend 651411) overnight. The next day blots were washed with TBST and incubated with HRP conjugated secondary antibodies. Lastly, western blotting luminol reagent solution was added to the membranes, and chemiluminescence signal was detected by Azure c600 Imaging Biosystems.

### Bone marrow-derived dendritic cell (BMDC) generation

Bone marrow was harvested from the femurs and tibia of an 8-week-old BALB/c mouse. Bones were flushed with a cold medium two times and centrifuged twice at 1500 rpm for 5 min. Cells were incubated with 20 ng/mL GMCSF (BioLegend 576304) at a density of 2 x 10^6^ cells for 8 days. On day 3, 20 ng/mL GMCSF in 10 mL was added, and on day 6, 10 mL of cell solution was replaced with 10 mL fresh medium containing 20 ng/mL GMCSF. On day 8 loosely adhering BMDCs were collected through gentle pipetting.

### Co-culture of spheroid conditioned medium with BMDCs

EMT6 spheroids were treated with [II] peptide, mitoxantrone, and cisplatin for 24h, and spheroid conditioned medium was mixed 1:1 (v:v) with BMDC medium, and BMDCs were incubated for 6 and 24h.

### Ethics

This study was carried out in accordance with the recommendations of the Guide for the Care and Use of Laboratory Animals from the National Institute of Health. Animal procedures were approved by the O.U. Health Sciences Center (OUHSC) Institutional Animal Care and Use Committee (protocol number R19-017A).

### In vivo

50.000 EMT6 cells were used to form a subcutaneous tumor model. Briefly, cells were injected in 100 μL of PBS and monitored for 8 days. At day 8, all the animals had palpable tumor formation, and the following treatment was done every two days: 0.5 mM [II] peptide in 0.2% MSA and 2% MSA in 100 μL of PBS. Tumor volume was measured every two days prior to injection and tumors were collected at day 22. Tumors were lysed using tip sonication and using RIPA Buffer (Thermo Scientific 89900) supplemented with protease and phosphatase inhibitor cocktail (Thermo Scientific 1861281). Protein concentrations were measured using BCA protein quantification kit (Thermo Scientific 23225). Cytokine concentrations were measured using ELISA and relative protein concentration was calculated through the following formula: Cytokine amount: (pg cytokine) / (mg tumor lysate).

### Enzyme-linked immunosorbent assay (ELISA)

Supernatants of BMDCs were collected after 6 and 24h incubation with spheroid conditioned mediums. Collected supernatants were centrifuged at 500g for 5 min at 4 ° C and used in ELISA. Tumor proteins were collected through tip sonication of the tumors at day 22. Supernatant was collected after centrifuging at 14.000g for 15 min. Briefly, plates were coated with capture antibody overnight. The next day, plates were blocked and incubated with supernatants, followed by incubation with the detection antibodies. Lastly, plates were incubated with Avidin-HRP and developed with TMB substrate solution. Absorbance was measured at 450 nm and 570 nm. 570 nm value was subtracted, and values were calculated based on the standard curve for IL-6 (BioLegend 431315), IL-1β (BioLegend 432615), IL-10 (BioLegend 431414), IFN-g (BioLegend 430804) and IL-12p70 (BioLegend 433604). Lipopolysaccharide (LPS) (Millipore Sigma L2630) was used as a positive control for cytokine release for BMDCs.

### Tissue Processing

Mouse kidney and liver tissue were collected and fixed in 4% PFA overnight, washed with PBS, and stored in 70% ethanol. Tissues were then dehydrated in increasing ethanol concentrations and cleared with multiple changes of xylene. The tissues were infiltrated by and embedded in paraffin wax using a Leica EG 1160 Paraffin Embedding Center and 5um sections were obtained by microtome (Microm HM325 Rotary Microtome). Samples were deparaffinized and rehydrated with changes of xylene and decreasing ethanol concentrations.

### Hematoxylin and Eosin (H&E) Stain

Regressive H&E staining was performed manually by transferring samples between plastic Coplin staining jars, proceeding in the following steps: staining with Hematoxylin Solution, Gill No. 3 (Sigma-Aldrich GHS332), differentiating with tap water and acidic ethanol, bluing with 0.1% sodium bicarbonate solution, staining with Eosin Y, 1%, Alcoholic Solution (EMS 26396-07), dehydrating, and clearing with xylene. Samples were mounted using Fisher Chemical™ Permount™ Mounting Medium (SP15-100) and VWR micro cover glass (VWR 48393-251) incubated overnight before imaging with Keyence BZ-X810 at 4x and 10x magnifications.

## 5 Acknowledgments

This work is partly supported by a grant from the Research Council of the University of Oklahoma Norman Campus and partly by the Oklahoma Tobacco Settlement Endowment Trust awarded to the University of Oklahoma, Stephenson Cancer Center. The content is solely the responsibility of the authors and does not necessarily represent the official views of the Oklahoma Tobacco Settlement Endowment Trust.

Research reported in this publication was supported in part by a Stephenson Cancer Center Pilot Grant (NIH -P30CA225520, funded by the National Cancer Institute Cancer Center Support Grant awarded to the University of Oklahoma Stephenson Cancer Center) and Oklahoma Center of Medical Imaging for Translational Cancer Research Pilot Grant (NIH - P20GM135009, funded by the National Institute of General Medical Sciences Support Grant awarded to the University of Oklahoma). The content is solely the responsibility of the authors and does not necessarily represent the official views of the National Institutes of Health.

